# The RNA helicase DDX21 cooperates with ETS1 and FLI1 in cell cycle, immune evasion, and snoRNA processing in activated B-cell-like diffuse large B-cell lymphoma cells

**DOI:** 10.64898/2026.06.02.729583

**Authors:** Giulio Sartori, Valdemar Priebe, Luciano Cascione, Mélanie Favre-Juilland, Vincenza Guzzardo, Marco Pizzi, Elaine Y.L. Chung, Sara Napoli, Alberto J. Arribas, Alessandro Ghiringhelli, Stefano Giuffrida, Mattia Forcato, Andrea Rinaldi, Paola Barraja, Silvio Bicciato, David W. Scott, Margot Thome, Francesco Bertoni

## Abstract

Diffuse large B-cell lymphoma (DLBCL) is a clinically and biologically heterogeneous disease, with the activated B-cell–like (ABC) subtype showing inferior outcomes. The ETS transcription factors ETS1 and FLI1 are recurrently gained and functionally relevant in DLBCL, yet their pathogenetic function is still to be fully elucidated. Here, we describe their cooperation with the RNA regulatory machinery, demonstrating that the RNA helicase DDX21 is a central effector of the ETS1/FLI1 transcriptional network in ABC-DLBCL. Our proteomic analyses revealed that ETS1 physically interacted with DDX21 and other RNA processing factors. As ETS1, DDX21 was preferentially expressed in ABC-DLBCL, particularly in the MCD/C5 genetic subtype, and it was associated with adverse clinical outcomes in this lymphoma subtype. Genetic and pharmacological studies demonstrated that DDX21 was essential for ABC-DLBCL cell proliferation. DDX21 also coordinated various transcriptional programs, which were revealed by integrated RNA-Seq, small RNA-Seq, ChIP-Seq, and Capture Hi-C analyses. DDX21-dependent transcription was relevant for ribosome biogenesis, MYC signaling, cell cycle progression, and immune evasion, but also regulated non-coding RNA networks, including microRNAs and small nucleolar RNAs, in particular SNORA37. Collectively, our data establish DDX21 as a nucleolar hub linking ETS transcription factors to coding and non-coding RNA programs that sustain aggressive lymphoma biology. These findings suggest DDX21 and ETS-RNA helicase complexes as promising therapeutic vulnerabilities in ABC-DLBCL.

## INTRODUCTION

Lymphomas are among the most common cancers and remain a leading cause of cancer deaths, especially in young adults ^1,2^. Diffuse large B-cell lymphoma (DLBCL) is the most common lymphoma, accounting for 30- 40% of cases, and is a clinically aggressive disease ^3^. Although many DLBCL patients can now be cured with chemoimmunotherapy regimens such as R-CHOP (rituximab, cyclophosphamide, doxorubicin, vincristine, and prednisone), approximately 30-40% develop refractory or relapsed disease, underscoring the need for improved therapeutic strategies ^3,4^. High-throughput genomic technologies have revealed that DLBCL comprises a heterogeneous group of biologically and clinically distinct entities ^5-7^. We have previously shown that the two ETS transcription factors, ETS1 and FLI1, co-mapped to the 11q24.3 region, are recurrently gained in up to 25% of DLBCL patients and co-regulate multiple genes involved in B-cell signaling, differentiation, and the cell cycle ^8-11^. Notably, FLI1 is more highly expressed in germinal center B-cell-like (GCB) DLBCL, whereas ETS1 predominates in the activated B-cell–like (ABC) subtype. The latter subtype is more aggressive and associated with inferior outcomes following standard therapy ^3^, highlighting the need to identify novel therapeutic vulnerabilities. In this study, by investigating the ETS1 interactome in ABC DLBCL, we identified the RNA helicase DDX21 as an additional therapeutic target functionally linked to ETS transcription factors.

## METHODS

### Cell lines

Cell lines were cultured under standard conditions at 37 °C in a humidified atmosphere with 5% CO_2_. Three ABC cell lines (HBL1, U2932, TMD8) and one GCB cell line (OCI-Ly1 and VAL) and were obtained and maintained as previously described ^12^. Cell line identity was confirmed by STR DNA fingerprinting. Mycoplasma contamination was excluded using the MycoAlert Mycoplasma Detection Kit (Lonza, Visp, Switzerland).

### Pull-down experiments and mass spectrometry

Protein precipitation was performed using streptavidin beads (2-1201-010, from IBA Lifesciences) to capture Strep-tagged ETS1 in the ABC DLBCL cell line HBL-1, followed by liquid chromatography-tandem mass spectrometry (LC-MS/MS). Shotgun proteomics analysis identified biologically relevant ETS1 protein interactors with spectral counts greater than four. Targets with a fold change > 4 in Strep-tagged ETS1 versus empty vector were identified using Scaffold software (Proteome Software, Inc., Portland, OR, USA), with a minimum peptide threshold of 90% and a protein threshold of 95%. Protein-protein association networks and functional enrichment analyses were performed with STRING ^13^.

### Immunoblots and co-immunoprecipitation experiments

For immunoblots, cells were harvested and lysed either by boiling samples in 2x Laemmli sample buffer (BioRad, Puchheim, Germany) supplemented with β-mercaptoethanol (Merck, Buchs, Switzerland) for 10′ or by following the manufacturer’s protocol using M-PER buffer (Thermo Fisher Scientific, Sede, Waltham, MA, USA). Lysates (30–50 μg) were resolved by electrophoresis on Mini-PROTEAN TGX Precast 4-20% gradient gels (BioRad, Puchheim, Germany) according to molecular weight. After electrophoresis, proteins were transferred to a nitrocellulose membrane (BioRad, Puchheim, Germany) by electrophoretic transfer, and the membranes were blocked in TBST (20 mM Tris-HCl [pH 7.5], 150 mM NaCl, 0.1% Tween 20) with 5% nonfat dry milk (BioRad, Puchheim, Germany) for 1 hour at room temperature (RT).

ETS1 protein interactors were validated by normal and reverse co-immunoprecipitation experiments. For each condition, 10 million cells were harvested, washed with PBS, and incubated on ice for 15 minutes with ice-cold lysis buffer (0.05% Triton, 100 mM NaCl, 20 mM Tris, pH 7.5, 1 mM EDTA, 1 mM EGTA) supplemented with Halt Protease and Phosphatase Inhibitor Cocktail (Thermo Fisher Scientific, Sede, Waltham, MA, USA). Lysates were cleared by centrifugation at 2200 × g for 15 minutes at 4°C. Lysates were incubated overnight at 4°C with 4 µg of the indicated primary antibody under gentle rotation. The following day, Protein A Magnetic Beads (Merck, Buchs, Switzerland) were added, and the mixture was incubated for 2 hours at 4°C. Beads were washed three times with ice-cold lysis buffer to remove nonspecifically bound proteins. Immunocomplexes were eluted in 2× SDS sample buffer and analyzed by immunoblotting with the indicated antibodies. Total cell lysates, inputs (2% of total protein used for IP) were run in parallel as controls. For immunocomplexes the following primary antibodies were used: rabbit polyclonal α-DDX21 (NB100-1718, Novus Biologicals), rabbit polyclonal α-SF3B1 (A300-996A, Bethyl Laboratories), rabbit polyclonal α-NOP56 (A302-721, Bethyl Laboratories), rabbit polyclonal α-DDX23 (ab70459, ABCAM). For immunoblotting the following primary antibodies were used in TBST with 5% BSA: rabbit polyclonal α-phospho-ETS1 (Thr38; ab59179, Abcam), mouse monoclonal α-SF3B1 (sc-514655, Santa Cruz Biotechnology), rabbit polyclonal α- NOP56 (A302-721, Bethyl Laboratories), mouse polyclonal α-THOC4 (ab6141, Abcam), mouse polyclonal α- DDX21 (sc-376953, Santa Cruz Biotechnology), rabbit polyclonal α-ETS1 (sc-350, Santa Cruz Biotechnology). Mouse monoclonal α-GAPDH (FF26A/F9, CNIO) was used in TBST with 5% nonfat dry milk. The secondary antibodies used were: ECL α-mouse IgG horseradish peroxidase-linked species-specific whole antibody and ECL α-Rabbit IgG horseradish peroxidase-linked species-specific whole antibody (GE Healthcare). When the same rabbit Ab was used for both IP and IB (ETS1 and NOP56) the Clean-Blot IP Detection Reagent (21230, Thermo Fisher Scientific) was used. Membranes were treated with Westar ηC 2.0 chemiluminescent substrate (Cyanagen, Bologna, Italy), and signals were detected using digital imaging with Fusion Solo (Vilber Lourmat, Lemont, IL, USA).

### RNA extraction, PCR amplification, quantitative real-time PCR

RNA was isolated using TRIzol (Invitrogen - Thermo Fisher, Waltham, MA, USA) and then DNase-treated with the RNase-free DNase Kit (Qiagen, Germantown, MD, USA). Total RNA extracts were reverse-transcribed using the SuperScript III First-Strand Synthesis SuperMix System (Invitrogen) to generate cDNA. In brief, 800 ng of total RNA was mixed with 10 μL of 2x RT Reaction Mix and 2 μL of RT Enzyme Mix, and the mixture was brought to a final volume of 20 μL with DEPC water (Invitrogen). Quantitative real-time (qRT)-PCR amplification was performed using the KAPA SYBR FAST qPCR Master Mix (2x) on an ABI Prism StepOnePlus Real-Time PCR system (Applied Biosystems). All primers were designed using the web-based program Primer3Plus (http://www.bioinformatics.nl/cgi-bin/primer3plus/primer3plus.cgi) and validated for target specificity with PrimerBlast (https://www.ncbi.nlm.nih.gov/tools/primer-blast/). The thermal cycler was programmed as follows: enzyme activation at 95 °C for 3′, followed by 40 cycles of denaturation (95 °C for 3 s) and annealing (60 °C for 30 s), and finally dissociation curve analysis. Primer efficiency was determined by linear modeling of the amplification curves with the LinReg software version 2015.4 ^14^. Relative quantification was calculated using the Pfaffl method ^15^. Primers targeting DDX21, SNORA37, and GAPDH are listed in Supplementary Table S1.

### Transcriptome analysis

Initial RNA quality control was performed on the Agilent BioAnalyzer (Agilent Technologies, CA, USA) using the RNA 6000 Nano kit (Agilent Technologies). RNA concentration was determined with the Invitrogen Qubit (Thermo Fisher Scientific) using RNA BR reagents (Thermo Fisher Scientific). Total RNA samples were prepared for RNA sequencing (RNA-Seq) with the NEBNext rRNA Depletion kit, the NEBNext Ultra Directional RNA Library Prep Kit for Illumina, and NEBNext Multiplex Oligos for Illumina (New England BioLabs Inc.). Sequencing was performed on a NextSeq 500 with the NextSeq 500/550 High Output Kit v2 (150 cycles PE; Illumina). All data are available in the National Center for Biotechnology Information (NCBI) Gene Expression Omnibus (GEO) database (http://www.ncbi.nlm.nih.gov/geo).

### Exon-intron split analysis (EISA)

To disentangle transcriptional from post-transcriptional regulatory effects, we performed Exon–Intron Split Analysis (EISA) using the eisa R package ^16^. Briefly, exon and intron read counts were quantified separately for each gene and sample and normalized by library size. Changes in intronic counts were treated as proxies for transcriptional regulation, whereas differences between exon and intron signals were used to infer post-transcriptional modulation. EISA scores and statistical tests were computed with default parameters unless otherwise stated.

### ChIP-Seq and ChIP qPCR

Direct targets were validated by DDX21 chromatin immunoprecipitation (ChIP). Chromatin was sheared with the M220 Focused ultrasonicator for Adaptive Focused Acoustics (AFA) technology (Covaris) using the milliTUBE 1 mL AFA fiber. The manufacturer’s protocol for the truCHIP Chromatin Shearing Kit was followed. 25 × 10^6 cells were washed in cold PBS and resuspended in Fixing Buffer A containing 1% formaldehyde and mixed for 2′. After crosslinking, the quenching buffer was added. Lysis of samples proceeded in accordance with the manufacturer’s protocol. The cell lysate suspension with chromatin was transferred into the milliTUBE and sonicated with the program set to 10% duty cycle with 200 cycles per burst for 12′. The quality of chromatin shearing was determined using the High Sensitivity DNA Analysis Kit (Agilent Technologies) and the 2100 BioAnalyzer (Agilent Technologies). ChIP was performed using 50 μL chromatin solutions (corresponding to 5 × 10^5 cells) diluted in ChIP dilution buffer (0.01% (w/v) SDS, 1.1% (v/v) Triton-X 100, 1.2 mM EDTA, 16.7 mM Tris–HCl, 167 mM NaCl [pH 8.1]) with 1x HALT Proteinase inhibitor cocktail (Thermo Scientific). Rabbit polyclonal α-DDX21 (5 μg, NB100-1718, Novus Biologicals),) was added to the diluted chromatin samples. Antibody/protein/DNA complexes were captured with protein A magnetic beads at 4 °C (Millipore). Magnetic beads were washed using a magnetic rack and increasing salt-stringency buffers in the following order: Low Salt washing buffer, 0.1% (w/v) SDS, 1% (v/v) Triton-X 100, 2 mM EDTA, 20 mM Tris– HCl [pH 8.1], 150 mM NaCl; High Salt washing buffer, 0.1% (w/v) SDS, 1% (v/v) Triton-X 100, 2 mM EDTA [pH 8.0], 20 mM Tris–HCl [pH 8.1], 500 mM NaCl; LiCl buffer, 0.25 M LiCl, 1% (w/v) IGEPAL-CA 630, 1% (v/v) deoxycholic acid, 1 mM EDTA, 10 mM Tris–HCl [pH 8.1]; Tris–EDTA buffer, 10 mM Tris–HCl, 1 mM EDTA [pH 8.1]. The Tris–EDTA buffer wash was repeated twice. Immunoprecipitated complexes were eluted from the dynabeads using elution buffer (1% (w/v) SDS, 0.1 M NaHCO3) with RNAse A added and incubated at 37 °C for 30′ on a thermomixer (1200 rpm). This was followed by reversal of cross-links by adding 5 M NaCl, 0.5 M EDTA, Tris–HCl, and Proteinase K for 2 h at 62 °C on a thermomixer (1200 rpm). Lastly, DNA was purified using the QIAquick PCR purification kit (Qiagen). For validation, qRT-PCR was performed using KAPA SYBR FAST qPCR Master Mix (2x) ABI Prism. Primers targeting DDX21, SNORA37, RPL23, and GAPDH are listed in Supplementary Table S11. For chromatin immunoprecipitation (ChIP) sequencing (ChIP-Seq), at least 5 parallel IPs were performed, and the eluted DNA was pulled and re-concentrated in 5 μL. All data will be available at the National Center for Biotechnology Information (NCBI) Gene Expression Omnibus (GEO) database (http://www.ncbi.nlm.nih.gov/geo).

### Data mining

All bioinformatic processing was performed using R/Bioconductor software packages in RStudio. RNA-Seq raw reads were quality assessed using fastqc ^17^. For each sample, the distribution of unique, multi-, and unmapped reads was checked for a high proportion of unmapped or multimapped reads. RNA sequencing reads were mapped against the human hg38 genome build using the Genecode version 22 annotation ^18^. Alignment was performed with STAR (v2.4.0 h) ^19^, and read counts overlapping gene features were obtained with HTSeq-Count ^20^. Transcripts with a count-per-million greater than one in at least three samples underwent differential gene expression analysis using the voom/limma ^21^ R package. Functional annotation was performed with g:Profiler using gene sets from the Molecular Signatures Database (MSigDB v5.1) ^22^ (Hallmark, c2.all, c5.bp, c6), SignatureDB ^23^, L1000 ^24^, Genomics of Drug Sensitivity in Cancer (GDSC) ^25^, and gene sets obtained from different publications as reported. Standard settings were used for g:Profiler data mining ^26^. Signatures with absolute log fold change > 0.1 and adjusted P < 0.05 were considered biologically relevant.

For ChIP-Seq analysis, raw sequencing reads were mapped to the Genome Reference Consortium Human Build 37 (GRCh37) using bowtie2. Read filtering was performed with SAMtools to retain reads that map uniquely and have a quality score of 10 or higher, and to remove duplicates. We first performed an exploratory analysis in the IGV genome browser to assess ChIP quality and detect issues and abnormalities. Peaks were then called using HOMER and selected to control the false discovery rate (FDR) at 0.001. To biologically interpret the results of ChIP-Seq experiments, we examined genes and other annotated elements in proximity to the identified enriched regions (peak annotations) using HOMER, PeakAnalyzer, and BedTools (version 2.17). Promoter regions were defined as within 3 kb of the closest TSS.

Target peaks located more than 3 kb from the closest TSS were annotated using the enhancer-promoter interactions map of Mifsud et al. ^27^, derived from a Capture HiC (C-HiC) experiment in GM12878 cells, a human Epstein-Barr virus (EBV)-transformed lymphoblastoid cell line. Active enhancers overlapping with target peaks were assigned to the corresponding interacting promoter region.

Pearson’s correlation was used to identify genes that were significantly (positively or negatively) correlated with DDX21 expression levels in DLBCL clinical specimens (GSE10846; p < 0.01). Overlapping between lists was done using the VENNY on-line tool ^28^.

### Data mining of publicly available datasets

Publicly available resources used included gene expression datasets for ETS1 and FLI1 ^10,11^; a B lymphoblasts ChIP-Seq dataset ^29^; RNA-Seq datasets for DLBCL (EGAD00001003783 ^30^, Centre for Lymphoid Cancer, Vancouver, BC, Canada; phs001444.v2.p1 ^31^, National Cancer Institute, Bethesda, MD, USA) and 3); and for mantle cell lymphoma (MCL) (EGAS00001004289 ^32^; Michael Smith Genome Sciences Centre, Vancouver, BC, Canada) clinical specimens. The CEL raw data files were imported and preprocessed using log2 transformation and normalization with the Bioconductor packages voom/limma ^21,33^ and edgeR ^34^ in RStudio ^35^. DDX21 mRNA expression was dichotomized into high and low values using the median as a cutoff for further analyses.

### Histological evaluation and immunohistochemical analysis

All cases were retrieved from the archives of the Surgical Pathology & Cytopathology Unit of Padua University Hospital (Padua, Italy). Each case was re-evaluated and assigned to a cell-of-origin subtype using the Hans algorithm ^36^. Immunohistochemical (IHC) analysis was performed on 4-μm-thick formalin-fixed, paraffin-embedded (FFPE) sections using the Bond Polymer Refine Detection kit on an automated immunostainer (BOND-MAX system; Leica Biosystems, Newcastle upon Tyne, UK), as previously described ^37^. The following primary antibodies were used: anti-CD10 (clone 56C6, Menarini Diagnostics, Florence, Italy), anti-Bcl6 (clone LN22, Leica Biosystems, Milan, Italy), anti-MUM1 (clone MUM1p, Dako, Glostrup, Denmark), and anti-DDX21 (HPA036593, Merck). Tissue microarrays were prepared as described previously ^38^, using the Galileo TMA CK3500 arrayer (Integrated System Engineering, Milan, Italy). Appropriate positive and negative controls were included. Phenotypic studies of benign palatine tonsils with reactive lymphoid hyperplasia were conducted in parallel to assess DDX21 expression in normal B cell subsets. DDX21 immunohistochemical expression was assessed as both (i) the percentage of positive neoplastic cells and (ii) the intensity of protein expression. The latter was defined by comparison with reactive germinal centers (GCs), as follows: score 0 = no expression; score 1+ = positive staining with intensity weaker than that of reactive GCs; score 2+ = positive staining with intensity equal to that of reactive GCs; score 3+ = positive staining with intensity stronger than that of reactive GCs.

### DDX21 gene silencing

For transient knockdown, we used the Amaxa 4D Nucleofector system (Lonza) to introduce four DDX21 siRNAs (J-011919-05, J-011919-06, J-011919-07, J-011919-08) from the ON-TARGET SMARTpool siRNA (L-011919) or a non-targeting siRNA as a control (Dharmacon GE Healthcare, now Horizon Discovery Ltd.). We followed the protocols for the SG Cell Line 4D-Nucleofector X Kit L (Lonza). In brief, 2 × 10^6^ cells were prepared and resuspended in 100 μL SG solution with 500 nM siRNA or the corresponding amount of BLOCK-iT (Invitrogen) as a control for nucleofection efficiency. Efficiency was confirmed 48 h after nucleofection by flow cytometry, and cells were harvested for protein lysates, RNA extraction, and an MTT [3-(4,5- dimethylthiazolyl-2)-2,5-diphenyltetrazoliumbromide] assay.

## RESULTS

### ETS1 directly interacts with DDX21, and other proteins involved in RNA processing

To identify ETS1 interaction partners in ABC-DLBCL, we expressed N- or C-terminally Strep-tagged ETS1 in HBL-1 cells, with an empty vector serving as a control. Pull-down experiments using streptavidin-coated beads were resolved by gel electrophoresis. Subsequently, immunoblots with ETS1 or pETS1 antibodies confirmed expression of Strep-tagged ETS1 and showed that the Strep-tag did not affect ETS1 phosphorylation at Thr38, regardless of its position (Figure S1A-B). Pull-down followed by LC-MS/MS identified 29 ETS1-binding proteins with spectral counts greater than 4 (0 in the empty vector) for either C- or N-terminally Strep-tagged ETS1 (Supplementary Table S1). Functional annotation revealed enrichment for proteins regulating RNA processing (Figure 1A), consistent with transcriptional signatures previously derived from ETS1 silencing ^9^ (Figure 1B). Immunoprecipitation confirmed interactions of ETS1 with DDX21, NOP56, THOC4 (ALYREF), and SF3B1 in HBL-1 cells expressing Strep-tagged ETS1 (Figure S2A) and in parental HBL-1 and U2932 cells expressing endogenous ETS1, and both derived from ABC DLBCL (Figure 1C and Figure S2B). In addition to direct co-immunoprecipitation using an ETS1 antibody, we performed reverse immunoprecipitation experiments with antibodies against target proteins in HBL-1 cells (Figure 1C); the binding of ETS1 to the two “spliceosome” proteins SF3B1 and THOC4 and to the two “ribosome” proteins NOP56 and DDX21 was reciprocally validated (Figure 1C). These findings were further confirmed in TMD8 (ABC DLBCL) and VAL (GCB DLBCL) cells (Figure 1D and Figure S2C). ETS1 phosphorylation at Thr38, which we previously showed to sustain DLBCL growth ^10^, did not alter ETS1-DDX21 interactions (Figure S2B). The resulting interaction network (Figure 1E) highlighted both known and novel ETS1-associated RNA processing factors.

**Figure 1.**
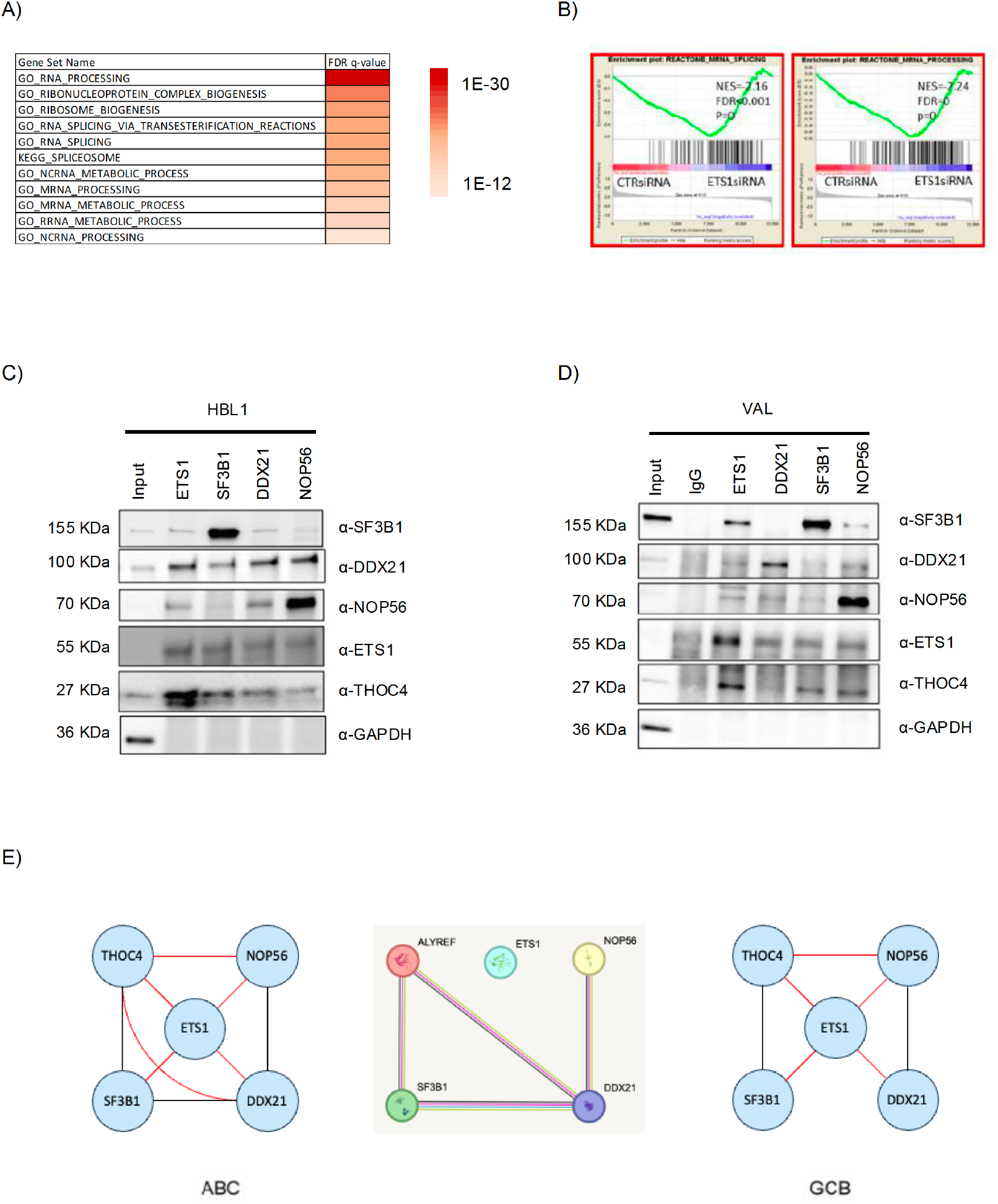
ETS1 regulates RNA processing by gene regulation and by protein-protein interaction. **(A)** Protein categories of potential ETS1 interactors identified by LS-MS/MS, with spectral count higher than 4, in either C- or N-terminally Strep-tagged ETS1 samples. FDR, false discovery rate. GO, gene ontology. (**B**) GSEA plots for mRNA splicing and mRNA processing that were enriched in ABC-DLBCL cell lines with the ETS1 knock down phenotype. (**C**) Immunoblot of ETS1 immunoprecipitates and input, and reverse co-immunoprecipitation in the ABC-DLBCL cell line HBL-1 (**D**) and the GCB-DLBCL cell line VAL. ETS1 interactions were confirmed by immunoprecipitation using antibodies for ETS1 or any of the indicated interactors. Positions of molecular weight markers are indicated. (**E**) Summary of co-immunoprecipitation results for ABC and GCB DLBCL. Black lines indicate known interactions while red lines indicate new interactions. The middle panel illustrates the multiple protein interactions identified by STRING; ALYREF is the official gene symbol for THOC4.

### DDX21 cooperates with ETS1/FLI1 and is essential in ABC-DLBCL

DDX21 is an RNA helicase involved in transcriptional regulation and ribosomal RNA processing ^39^, expressed in human marginal zone B cells ^40^, and overexpressed in several human cancers, including lymphomas ^41-43^. Its functional role in DLBCL remains unexplored. Given the potential therapeutic implications, as we and others have reported using small molecules that disrupt the binding of ETS factors and RNA helicases ^44,45^, we decided to study this helicase in ABC DLBCL. Analysis of available gene expression and ChIP-Seq datasets ^10,11,29^ showed that DDX21 is a transcriptional target of both ETS1 and FLI1 in B cells (data not shown). We therefore assessed DDX21 levels in a previously established model in which we genetically inhibited IgM stimulation-induced phosphorylation of ETS1 at threonine 38, a marker of ETS1 activation ^10^. DDX21 levels were lower in mutant pETS1 samples (vs untreated) (logFC=0.26, adj.P.=0.009) than in pETS1 wild-type samples (vs untreated) (logFC=0.33, adj.P.<0.001). Furthermore, DDX21 appeared to be an ETS1 target gene in human B lymphoblasts (data not shown).

Next, we validated FLI1 binding at the DDX21 promoter in HBL-1 and U2932 cells (Figure 2A) and confirmed DDX21 downregulation upon FLI1 or DDX21 silencing (Figure 2B). DDX21-bound regions were enriched for ETS1/FLI1 motifs (Suppl. Figure 3A) and for pathways linked to MYC, RNA processing, proliferation, DNA repair, and apoptosis (Suppl. Figure 3B).

**Figure 2.**
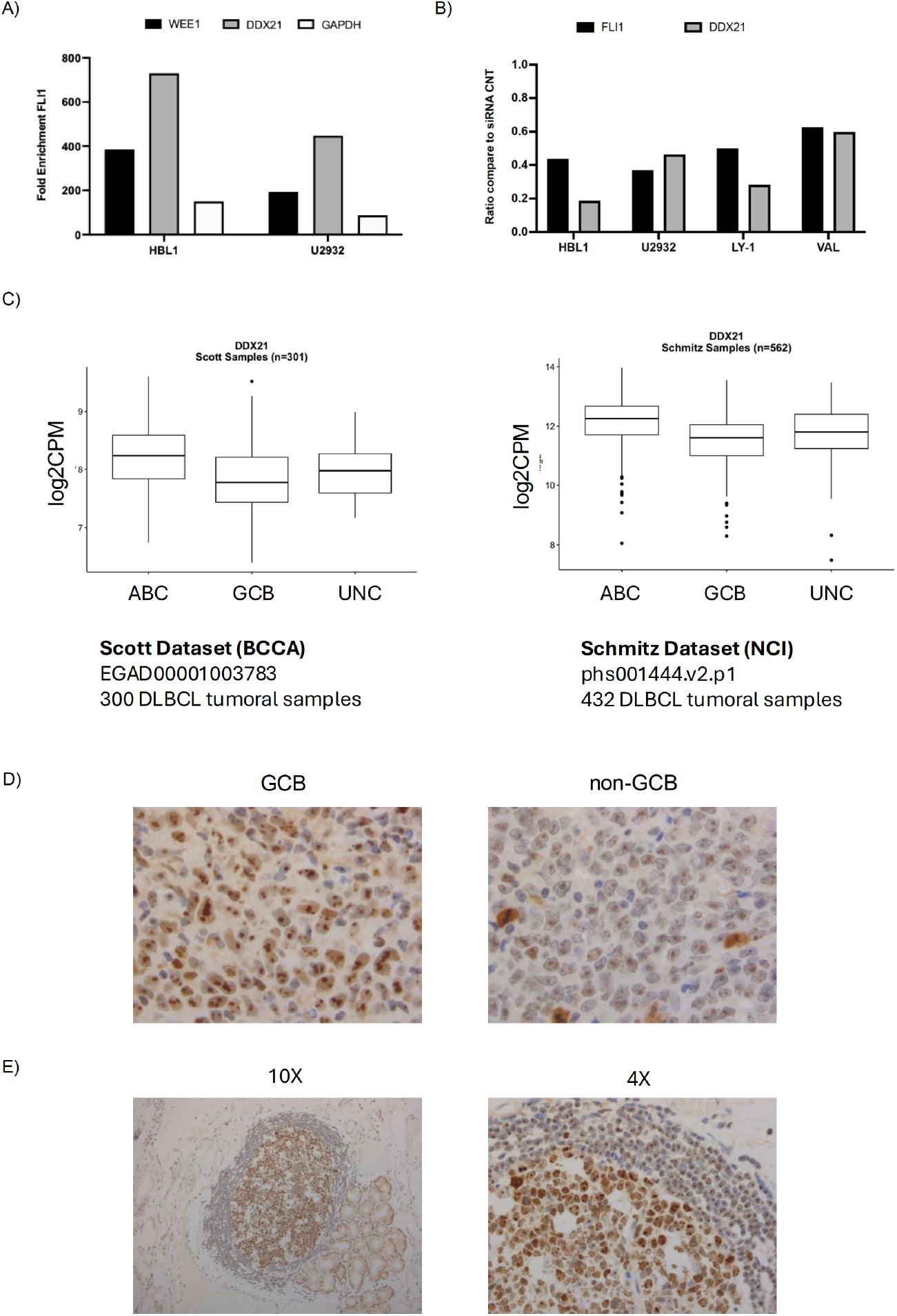
DDX21 is regulated by FLI1, and it is an important protein in ABC DLBCL cell line. **(A)** Validation by qRT-PCR of ChIP-Seq data showing FLI1 binding to DDX21 promoter regions. The figure shows the relative quantification of promoter regions bound by FLI1 in in two different ABC-DLBCL cell lines, HBL1 and U2932. WEE1 and GAPDH amplification were used as a positive and negative control, respectively. **(B)** Relative mRNA expression (normalized to GAPDH) of FLI1 and DDX21 from control CNT siRNA and FLI1 siRNA treated cells. **(C)** Differential expression of FLI1 RNA in two datasets (EGAD00001003783 and phs001444.v2.p1) comparing GCB DLBCL to ABC DLBCL. UNC: unclassified DLBCL. (**D**) Immunohistochemistry staining showing DDX21 in DLBCL clinical specimens, with higher intensity in non-GCB (left) compared to GCB (right) in two representative DLBCL clinical specimens. (**E**) Immunohistochemistry staining of a reactive germinal center at 10X (left) and 40X (right).

**Figure 3.**
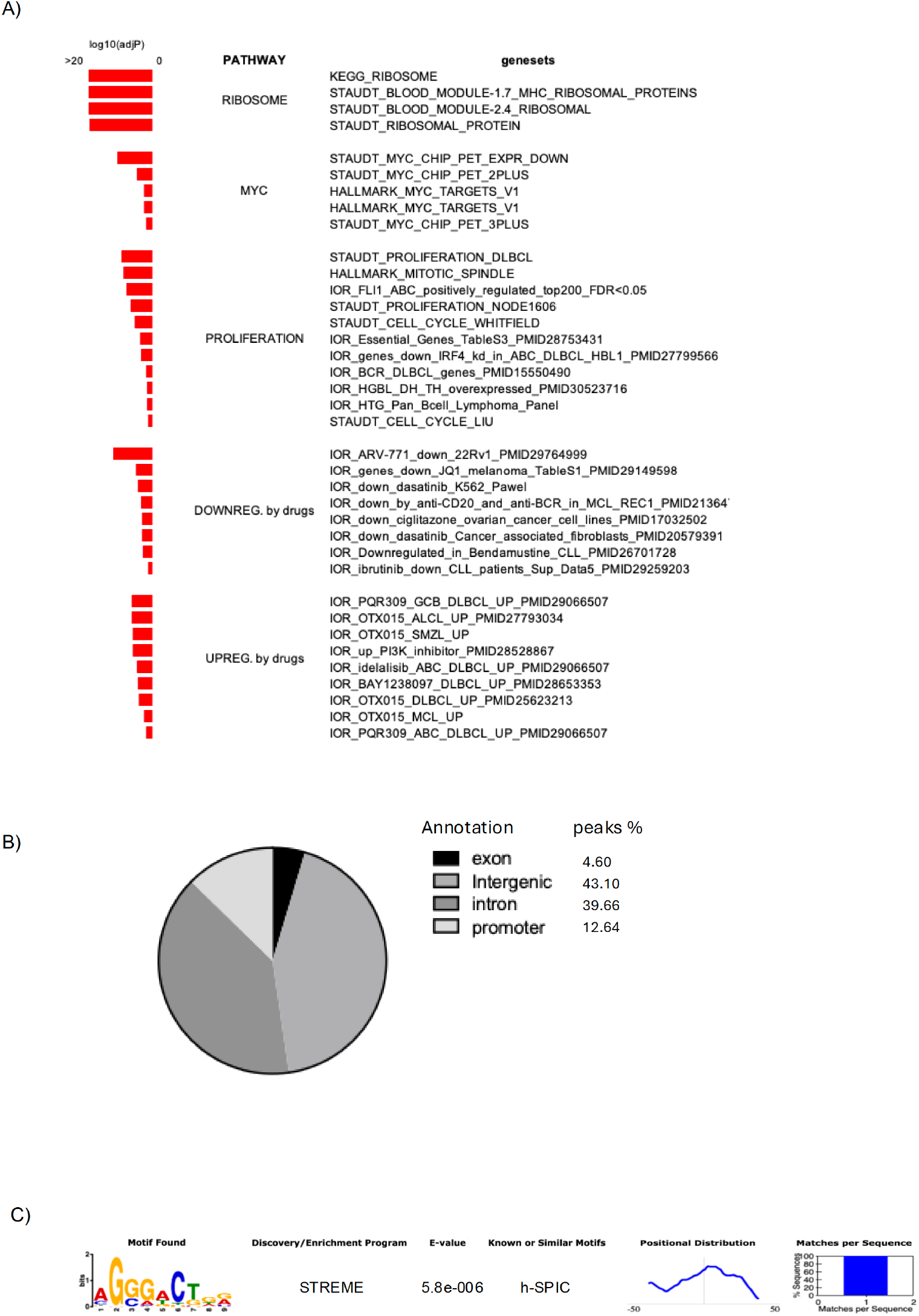
Pathway analysis following DDX21 downregulation and genomic distribution of FLI1 binding sites identified by ChIP-Seq. **(A)** Summary of pathways enriched in DDX21 up- (red) or downregulated (blue) genes after RNA-Seq, comparing DDX21 knockdown versus control samples in the ABC DLBCL cell lines HBL1 and U2932 (absolute logfold change > 0.1 and adj.P < 0.05). Significant g:GOSt annotated pathways/signatures (adjusted p-value < 0.05) are grouped into biological processes and sorted by adjusted p-value. (**B)** Distribution of DDX21 binding sites as assessed by ChIP-Seq in both ABC DLBCL cell lines. (**C)** ETS-like consensus binding motif enrichment found with MEME among ChIP peaks.

In clinical datasets, DDX21 RNA expression correlated positively with ETS1, FLI1, and their interactors (Supplementary Table S2) and was significantly higher in ABC than in GCB DLBCL (Figure 2C). Consistent with these results, we observed higher DDX21 expression in the MCD group than in all other genetic subgroups (Suppl. Figure 3C). Immunohistochemistry confirmed stronger DDX21 staining in non-GCB than in GCB cases (Figure 2D; Supplementary Table S3) and showed staining in reactive germinal centers (Figure 2E). High DDX21 expression was associated with poor outcomes in ABC DLBCL but not in GCB DLBCL cohorts (Suppl. Figure 4A–B). Finally, DDX21 silencing reduced proliferation in ABC cell lines (Suppl. Figure 4C–D) as confirmed by the dependency scores calculated using published CRISPR data ^31^ (Suppl. Figure 4E). Taken together, these results highlight an important role for DDX21 as a FLI1- and ETS1-induced gene whose expression drives proliferation in ABC DLBCL.

**Figure 4.**
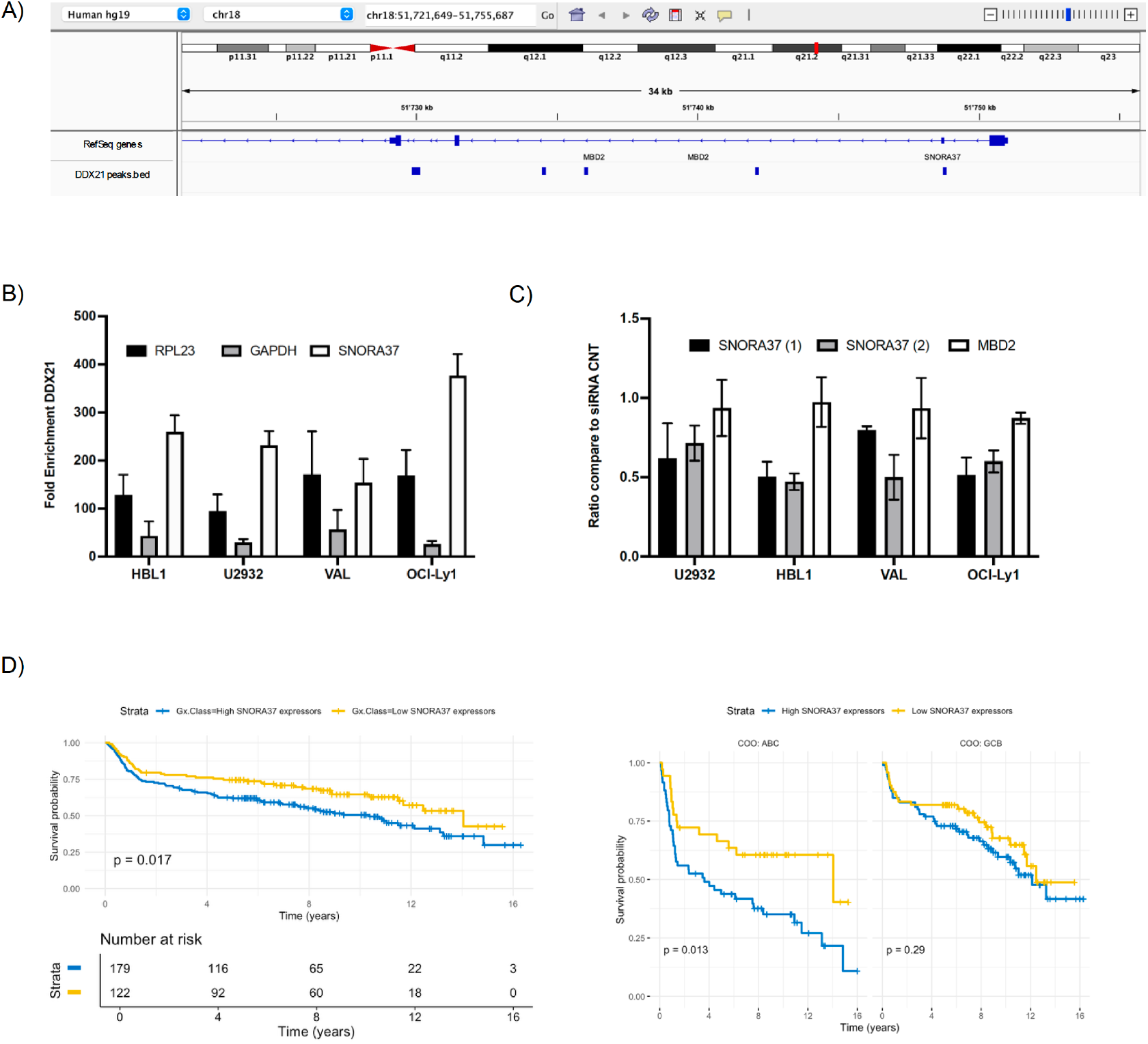
SNORA37 is a direct transcriptional target of DDX21 with a prognostic role in ABC DLBCL. **(A)** DDX21 binding in the promoter region of SNORA37 as assessed by ChIP-Seq **(B)** Validation by qRT-PCR of ChIP-Seq data showing DDX21 binding in the SNORA37 promoter region. The figure shows the relative quantification of DDX21-bound promoter regions in HBL1 and U2932. GAPDH amplification was used as a negative control. (**C**) Relative mRNA expression (normalized to GAPDH) of SNORA37 in DDX21 siRNA treated cells. MBD2 is SNORA37 host gene. (**D**) Overall Survival (OS) Kaplan-Meier curves in the cohort of patients dichotomized by SNORA37 expression level, with 2 counts per million (cpm) as the cutoff, yielding two classes: “*High SNORA37 expressors*” (blue) and “*Low SNORA37 expressors*” (yellow). Raw data obtained from CLC (EGAD00001003783) (X-axis, years; Y-axis, percentage of patients).

### DDX21 orchestrates transcriptional and non-coding RNA programs in ABC-DLBCL

Immunohistochemistry of DLBCL clinical specimens showed that DDX21 localized predominantly to nucleoli in all analyzed DLBCL biopsies (Fig. 2D), consistent with its role in transcriptional control and ribosomal RNA metabolism. Therefore, we next analyzed the broader effects of siRNA-mediated DDX21 silencing on gene transcription. RNA-Seq following DDX21 knockdown in ABC DLBCL cells (Suppl. Figure 5A-C) revealed that DDX21 positively regulated gene programs enriched for ribosome biogenesis, MYC targets, and pathways sensitive to BTK, PI3K/mTOR, and BET inhibitors (Figure 3A and Suppl. Figure 5D). DDX21-driven transcripts included *AICDA, ERBB4, HDAC3/11, MCM8*, and *CD274* (PD-L1), whereas those negatively regulated by the helicase were enriched for IL6/IL10 signaling and MHC class I and II components (Supplementary Table S4), suggesting a contribution to immune evasion. We then used the EISA computational approach ^16^, which quantifies changes in mature RNA and pre-mRNA reads across experimental conditions to assess transcriptional and post-transcriptional regulation of gene expression. This revealed that DDX21 primarily acted at the transcriptional level rather than post-transcriptionally. Indeed, post-transcriptionally modulated genes were present but were discordant across cell lines and did not correlate with the logFC of DDX21 (Suppl.Figure 5E). Small RNA-Seq after DDX21 silencing (Supplementary Table S5) uncovered a complementary layer of regulation involving microRNAs (miRNAs), small nuclear RNAs (snRNAs), and small nucleolar RNAs (snoRNAs). DDX21 depletion upregulated let-7 family members and downregulated miR-222 and miR-155, consistent with its pro-oncogenic transcriptional effects. Several snRNAs and snoRNAs were also altered, among which SNORA37 emerged as strongly upregulated. These results suggested that DDX21 coordinates both coding and non-coding RNA networks from its nucleolar hub to sustain the growth of ABC-DLBCL.

### DDX21 directly controls chromatin-associated transcription and SNORA37 expression

Next, we performed ChIP-Seq analysis, integrating our results with available Capture Hi-C (C-HiC) data ^27^, followed by immunoblotting and qRT-PCR (Suppl. Figure 6). We identified over 4,900 DDX21 binding peaks, predominantly distal to promoters (Figure 3B and Supplementary Table S6), and linked them to 1,225 candidate target genes via C-HiC analysis (Supplementary Table S6). Motif enrichment revealed ETS-like (Figure 3C), PAX5, FOS, and IKZF1 motifs (Suppl. Fig. 6), consistent with co-regulatory functions. Integration of RNA-Seq with additional publicly available ChIP ^39^ and CRISPR screens ^46^ highlighted DDX21-dependent regulation of ribosomal genes (e.g., *RPL3, RPS2*), metabolic genes (e.g., *PGAM1*), and therapy resistance– associated genes (e.g., *PABPC1*. Integration of ChIP-Seq, C-HiC, and small RNA-Seq data identified a small nucleolar RNA with oncogenic activity ^47^, SNORA37, as a direct transcriptional target of DDX21, with DDX21 binding detected at both promoter-proximal and distal enhancer regions). The corresponding ChIP peak (Figure 4A) was validated by ChIP-qPCR (Figure 4B), and SNORA37 expression was specifically downregulated upon DDX21 silencing (Figure 4C) in ABC and GCB DLBCL cell lines. SNORA37 was identified as a prognostic biomarker in the previously analyzed ABC DLBCL cohort (Figure 4D), whereas its host gene, MBD2, was not prognostic in DLBCL. Collectively, these findings demonstrate that DDX21 directly drives SNORA37 expression in ABC DLBCL, consistent with its known pro-oncogenic effects ^48,49^.

## DISCUSSION

By integrating proteomic, genomic, and functional approaches, we established DDX21, an RNA helicase involved in transcriptional regulation and ribosomal RNA processing, as a critical effector of the ETS1/FLI1 transcriptional network in ABC DLBCL. DDX21 not only physically interacted with ETS1 but was also transcriptionally regulated by both ETS1 and FLI1, thereby creating a reinforcing circuit that sustains the gene expression programs of the malignant B cell. These programs included ribosome biogenesis, MYC signaling, and pathways linked to immune evasion, underscoring DDX21’s role in lymphoma biology.

Previous studies have shown that DDX21 sustains ribosome biogenesis and participates in resolving R-loops and G-quadruplexes, processes critical for genome stability ^39,50-52^. DDX21 is also recruited by ADAR1 alongside DHX9 to unwind R-loops, linking RNA helicases to transcription termination and DNA repair ^53,54^. Additional evidence shows that DDX21 localizes to the nucleolus, where it occupies the transcribed rDNA locus, directly contacts both rRNA and snoRNAs, and promotes rRNA transcription ^39,55^. DDX21 promotes tumorigenesis in colorectal cancer ^48^ and predicts survival outcomes in gynecologic cancers ^49^, similar to what we showed here in DLBCL.

ETS1 and FLI1, recurrently gained in DLBCL, have long been associated with aberrant transcriptional control ^8-11^. Our findings expand this network by incorporating DDX21 and other RNA processing proteins, suggesting a cooperative mechanism that promotes ABC DLBCL aggressiveness. Convergence at both protein and DNA levels positions DDX21 as a nexus connecting transcription factors with RNA metabolism. DDX21 was preferentially expressed in ABC compared to GCB DLBCL, with the highest expression in the MCD genetic subtype, and its expression correlated with adverse prognosis in both DLBCL and MCL, another NF-κB-driven lymphoma. DDX21 was also essential for lymphoma cell survival, consistent with previously reported large-scale genetic screens ^46,56^.

Transcriptomic analyses revealed that DDX21 activates MYC targets and survival pathways while repressing immune signaling and antigen presentation. These dual roles highlight DDX21 as both an oncogenic driver and an immune-evasion mediator. RPL and RPS genes were confirmed as direct targets of DDX21, as previously reported ^39^. In addition, PABPC1, which regulates therapy resistance in chronic myeloid leukemia ^57^, and PGAM1, whose inhibition promotes ferroptosis and synergizes with anti-PD-1 immunotherapy in hepatocellular carcinoma models ^58^, were also identified. Our ChIP-Seq analysis further showed that DDX21 binds to genomic regions enriched for ETS, PAX5, FOS, and IKZF1 motifs, consistent with previous reports that DDX21 cooperates with other transcription factors to regulate transcription ^39,59^.

We uncovered a role for DDX21 in regulating non-coding RNAs. Small RNA-Seq identified deregulated miRNAs, including oncogenic miR-155 and miR-222 ^60,61^, as well as snoRNAs such as SNORA37, recently implicated in gastric cancer progression ^47^. Consistent with our data, SNORA37 was found as a target gene in ChIP data of ETS1 (GSM803510 with a score of 0.901) and DDX21 (GSM1369392 with a score of 0.555) while not in FLI1 (GSM1097880 with a score of 0.092), and, based on preferential association of DDX21 with rRNA modification sites, snoRNAs represent the principal class of DDX21-bound RNAs ^39^. *In vitro* experiments demonstrated that several snoRNAs, including SNORA37, directly bind to the DNA-binding domain of PARP-1, thereby increasing PARylation activity. This snoRNA-mediated autoPARylation of PARP-1 further potentiates its interaction with the DEAD-box RNA helicase DDX21, thereby promoting rDNA transcription ^62^. This extends the functional scope of DDX21 to small RNA networks, which are increasingly recognized as contributors to lymphoma biology ^55,62^.

Our findings suggest that DDX21-snoRNA circuits may represent a novel oncogenic dependency. These insights have therapeutic implications. RNA helicases are emerging as druggable vulnerabilities ^63,64^. Small molecules such as YK-4-279 and TK-216 disrupt ETS-RNA helicase interactions and show anti-lymphoma activity ^44,45^, and they also affect the expression of various snoRNAs ^45,65^. Our results, supported by genetic and pharmacological inhibition of DDX21, provide a rationale for exploring similar strategies to inhibit ETS1- DDX21 complexes or DDX21 itself. Given its integration of MYC signaling, ribosome biogenesis, and immune evasion, targeting DDX21 may synergize with BTK/PI3K, immune checkpoint inhibitors, or epigenetic agents. A lead compound, KI-DX-014, that inhibits the interaction between DDX21 and RNA, thereby modulating the RNA-dependent functions of DDX21, has been recently described, but its poor membrane permeability ^64^ does not yet allow its testing in cellular models ^64^.

In conclusion, DDX21 emerges as a central regulator in ABC DLBCL by bridging ETS transcription factors with transcriptional and post-transcriptional programs. Through its impact on both coding and non-coding RNAs, DDX21 reinforces oncogenic pathways and promotes immune evasion, making it a promising therapeutic target in aggressive B-cell lymphomas.

## Acknowledgment

This research was funded by the Swiss National Science Foundation (Sinergia grant number CRSII3_147620/1, to FB and MT), and by Rotary Foundation grants GG1639200 and GG1756935 (to GS). MT acknowledges support from Swiss Cancer Research (KFS-5411-08-2021) and the Emma Muschamp Foundation. The authors used OpenAI’s ChatGPT (GPT-5.4, OpenAI, San Francisco, CA, USA) to assist in text preparation and drafting. After using this tool/service, the authors reviewed and edited the content as needed and take full responsibility for the content of the published article.

