## Supplementary materials for "The RNA helicase DDX21 cooperates with ETS1 and FLI1 in cell cycle, immune evasion, and snoRNA processing in activated B-cell-like diffuse large B-cell lymphoma cells"

### Supplementary Figures

**Figure S1. Expression and phosphorylation of Strep-tagged ETS1 constructs.** (**A**) Immunoblot showing ETS1 expression in lysates or immunoprecipitations obtained from HBL-1 cells. (**B**) Immunoblot assessing ETS1 Thr38 phosphorylation in lysates of HBL-1 cells expressing Strep-tagged ETS1 or immunoprecipitation samples. ETS1-Strep has an expected molecular weight of 59 kDa, and Strep-ETS1 of 57 kDa. Endogenous ETS1 is expected to migrate at 55 kDa. Immunoprecipitation was done using α-Strep antibody. Positions of molecular weight markers are indicated.

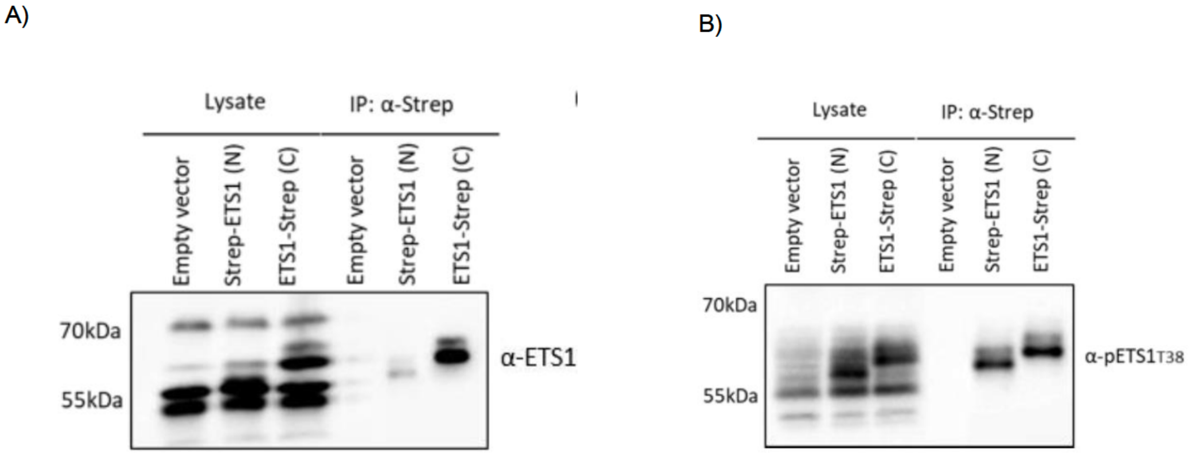

**Figure S2. Validation of protein interactions by co-immunoprecipitation.** (**A**) Immunoblot of input, control, and ETS1 immunoprecipitates from the ABC-DLBCL cell line HBL-1 (parental or expressing Strep-tagged ETS1, which migrates just above endogenous ETS1). Samples were analyzed by immunoblot using the indicated antibodies. (**B**) Immunoblot of total cell lysates, inputs, controls, and ETS1 immunoprecipitates from the ABC-DLBCL cell line U2932 after IgM stimulation for 20 minutes. (**C**) Immunoblot of total cell lysate, input, and reverse co-immunoprecipitations from the ABC-DLBCL cell line TMD8. ETS1 interactions were confirmed by immunoprecipitation using antibodies against ETS1 or any of the indicated interactors. Positions of molecular weight markers are indicated.

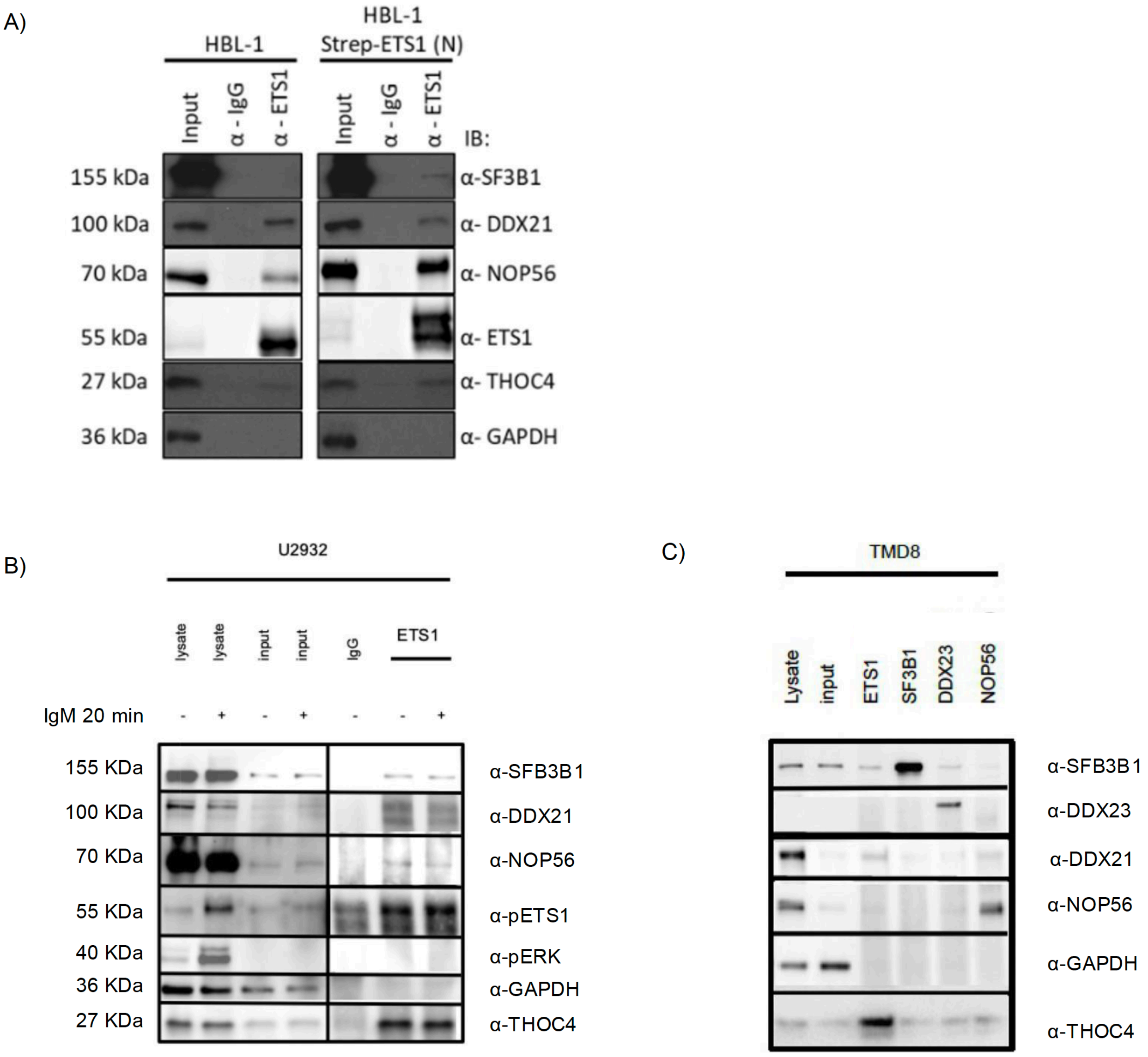

**Figure S3. DDX21-bound DNA regions are enriched in ETS1/FLI1 recognition motifs.** (**A**) Overlapping of FLI1, ETS1, and DDX21 targets. (**B**) Annotation of peaks common to the three transcriptional factors. (**C**) Differential expression of DDX21 RNA in GSE117556 and phs001444.v2.p1 based on genetic subtypes. (**D**) DDX21 downregulation in DLBCL cell lines harvested 72 h after nucleofection. (**E**) MTT assay for DLBCL cell lines nucleofected with either 500 nM control (CNT) siRNA or DDX21 siRNA. **(F)** Cumulative plot showing the dependency scores calculated using data from 8 cell lines [1]. A negative score indicates that the gene is more likely to be essential in DLBCL cell lines.

A)

**FLI1**: GSM1097880
Cell line: CD34+
Cell type: Haematopoietic Progenitor Cell

**FLI1:** GSE157191
Cell line: (2 GCB cell lines) plus 2 ABC cell lines
Cell type: DLBCL

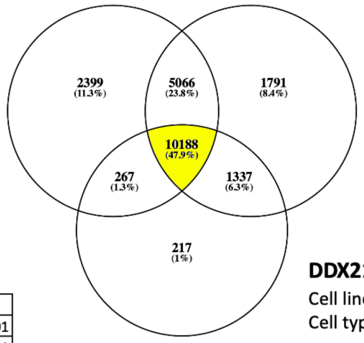


**ETS1:** GSM803510
Cell line: GM12878
Cell type: B Lymphocyte

**DDX21:** GSM1369392
Cell line: HEK293
Cell type: Epithelium

B)

| Gene set name | p.Val |
|---|---|
| HALLMARK_MYC_TARGETS_V1 | <0.001 |
| KEGG_RIBOSOME | <0.001 |
| KEGG_SPLICEOSOME | <0.001 |
| HALLMARK_DNA_REPAIR | <0.001 |
| HALLMARK_E2F_TARGETS | <0.001 |
| HALLMARK_MYC_TARGETS_V2 | <0.001 |
| KEGG_OXIDATIVE_PHOSPHORYLATION | <0.001 |
| HALLMARK_P53_PATHWAY | 0.001 |
| HALLMARK_PI3K_AKT_MTOR_SIGNALING | 0.002 |

C)

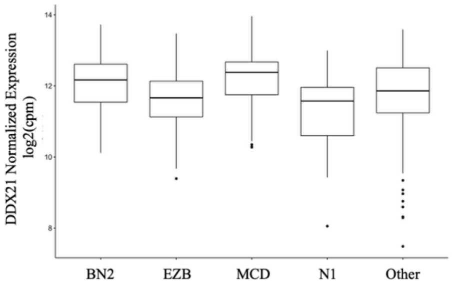


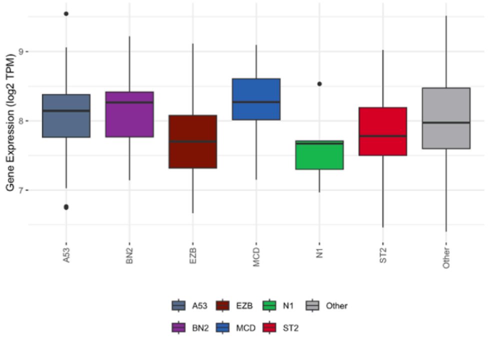


**Schmitz Dataset (NCI)**
phs001444.v2.p1
432 DLBCL tumoral samples

**Scott Dataset (BCCA)**
EGAD00001003783
300 DLBCL tumoral samples

**Figure S4. DDX21 expression is associated with inferior outcome in DLBCL patient's dataset.** Overall Survival (OS) Kaplan-Meier curves in the cohort of patients dichotomized using DDX21 expression level, setting 2 counts per million (cpm) as cutoff, obtaining two classes: "*High DDX21 expressors*" (blue) and "*Low DDX21 expressors*" (yellow) in (**A**) DLBCL patients (**B**) ABC DLBCL and (**C**) GCB DLBCL patients. Raw data obtained from CLC (EGAD00001003783) (X-axis, years; Y-axis, percentage of patients). (**D**)

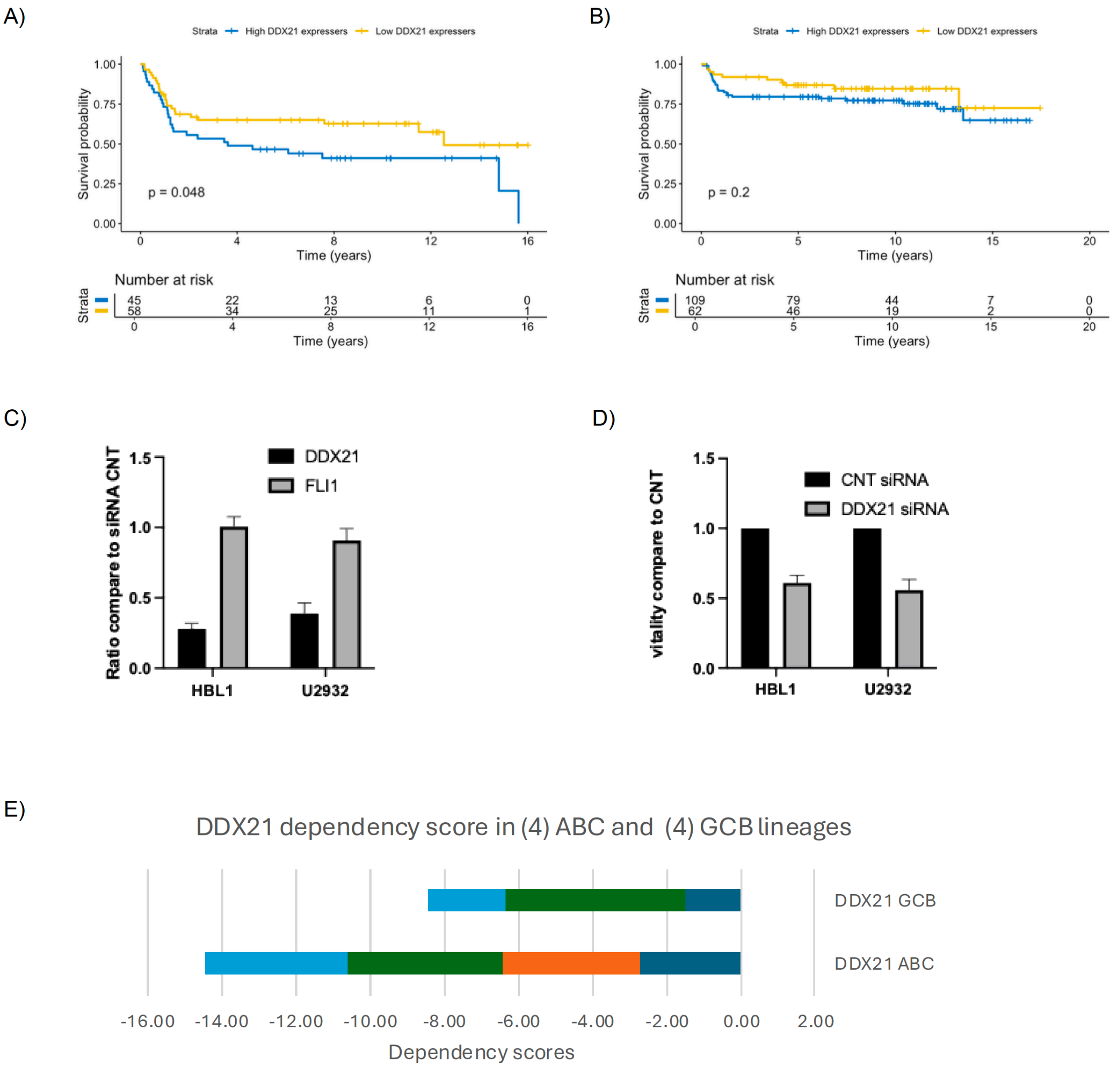

**Figure S5. DDX21 downregulation in ABC DLBCL cell lines harvested 48 hrs after nucleofection with either 500nM CNT siRNA or DDX21 siRNA. (A**) Normalized (to GAPDH) relative mRNA expression of DDX21 from CNT siRNA and DDX21 siRNA-treated cells. (**B**) Immunoblot showing DDX21 protein expression in CNT siRNA- and FLI1 siRNA-treated cells. Mouse monoclonal α-GAPDH was used as a loading control. Three different replicates are shown. (**C**) GSEA plots for gene expression signatures were obtained. Green line, enrichment score; bars in the middle portion of the plots show where the members of the gene set appear in the ranked list of genes; Positive or negative ranking metric indicates, respectively DDX21 upregulated and downregulated genesets; FDR, false discovery rate; NES, normalized enrichment score.

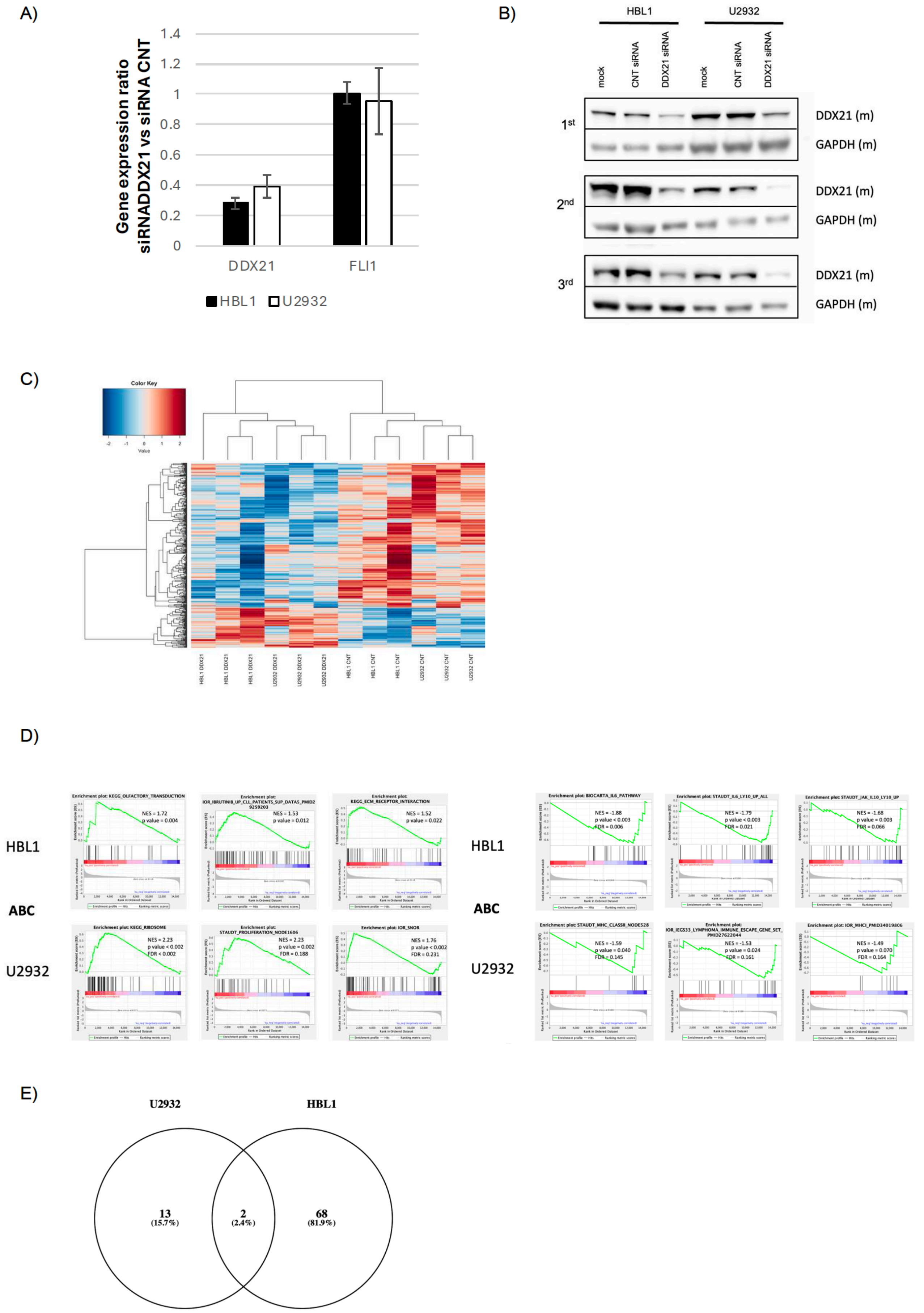
A)
Gene expression ratio siRNADDX21 vs siRNA CNT
1.4
1.2
1
0.8
0.6
0.4
0.2
0
DDX21
FLI1
HBL1
U2932
B)
HBL1
U2932
mock
CNT siRNA
DDX21 siRNA
mock
CNT siRNA
DDX21 siRNA
1st
2nd
3rd
DDX21 (m)
GAPDH (m)
DDX21 (m)
GAPDH (m)
DDX21 (m)
GAPDH (m)
C)
Color Key
Value
HBL1 DDX21
HBL1 DDX21
HBL1 DDX21
U2932 DDX21
U2932 DDX21
U2932 DDX21
HBL1 CNT
HBL1 CNT
HBL1 CNT
U2932 CNT
U2932 CNT
U2932 CNT
D)
HBL1
ABC
U2932
Enrichment plot: KEGG_OLFACTORY_TRANSDUCTION
NES = 1.72
p value = 0.004
NES = 1.53
p value = 0.012
Enrichment plot: KEGG_ECM_RECEPTOR_INTERACTION
NES = 1.52
p value = 0.022
Enrichment plot: KEGG_RIBOSOME
NES = 2.23
p value < 0.002
FDR < 0.002
Enrichment plot: STAUDT_PROLIFERATION_NODE1606
NES = 2.23
p value = 0.002
FDR = 0.188
Enrichment plot: IOR_SNOR
NES = 1.76
p value < 0.002
FDR = 0.231
HBL1
ABC
U2932
Enrichment plot: BIOCARTA_IL6_PATHWAY
NES = -1.88
p value < 0.003
FDR = 0.006
Enrichment plot: STAUDT_IL6_LY10_UP_ALL
NES = -1.79
p value < 0.003
FDR = 0.021
Enrichment plot: STAUDT_JAK_IL10_LY10_UP
NES = -1.68
p value = 0.003
FDR = 0.066
Enrichment plot: STAUDT_MHC_CLASSII_NODE528
NES = -1.59
p value = 0.040
FDR = 0.145
NES = -1.53
p value = 0.024
FDR = 0.161
Enrichment plot: IOR_MHCI_PMID34019806
NES = -1.49
p value = 0.070
FDR = 0.164
E)
U2932
HBL1
13
(15.7%)
2
(2.4%)
68
(81.9%)

**Figure S6. Validation by immunoblot and real-time PCR of ChIP-Seq experiments data.** A) Immunoblot showing presence of DDX21 and GAPDH in DDX21 immunoprecipitates obtained from the indicated ABC and GCB DLBCL cell lines. B) Relative quantification of promoter regions bound by DDX21 in HBL1 and U2932. RPL23 amplification was used as a positive control while GAPDH as a negative control. C) Consensus binding motifs enrichment found with MEME among ChIP peaks.

A)

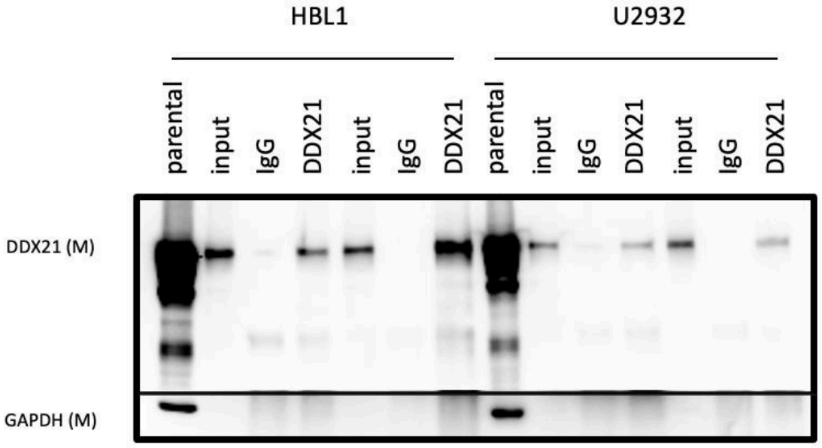
HBL1
U2932
parental
input
IgG
DDX21
input
IgG
DDX21
parental
input
IgG
DDX21
input
IgG
DDX21
DDX21 (M)
GAPDH (M)

B)

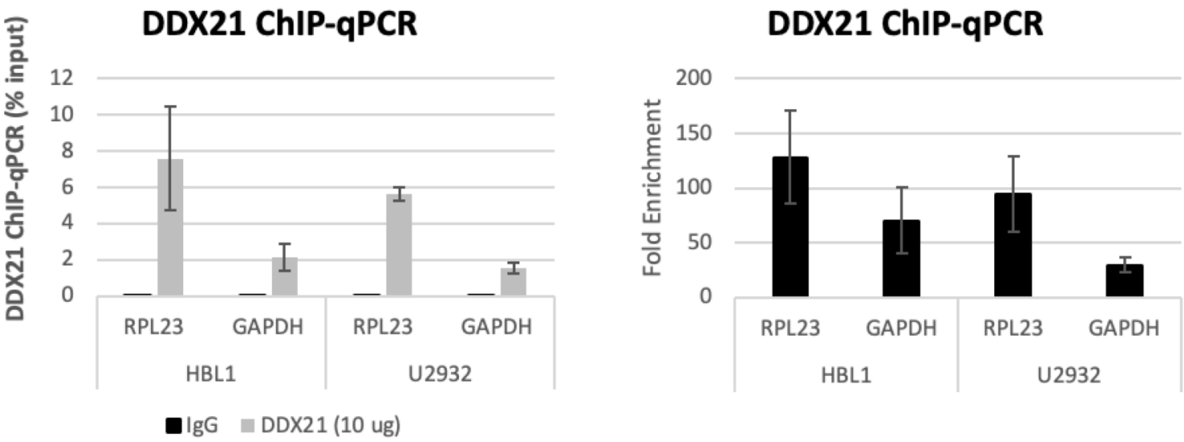
DDX21 ChIP-qPCR
DDX21 ChIP-qPCR (% input)
12
10
8
6
4
2
0
RPL23
GAPDH
RPL23
GAPDH
HBL1
U2932
IgG
DDX21 (10 ug)
DDX21 ChIP-qPCR
Fold Enrichment
200
150
100
50
0
RPL23
GAPDH
RPL23
GAPDH
HBL1
U2932

C)

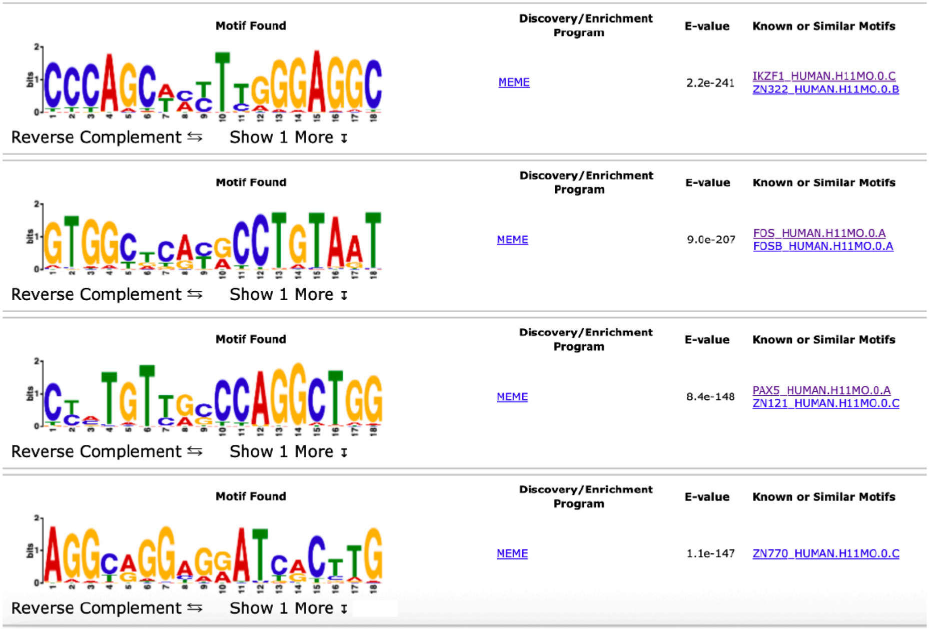
Motif Found
Discovery/Enrichment Program
E-value
Known or Similar Motifs
MEME
2.2e-241
IKZF1_HUMAN.H11MO.0.C
ZN322_HUMAN.H11MO.0.B
Reverse Complement
Show 1 More
Motif Found
Discovery/Enrichment Program
E-value
Known or Similar Motifs
MEME
9.0e-207
FOS_HUMAN.H11MO.0.A
FOSB_HUMAN.H11MO.0.A
Reverse Complement
Show 1 More
Motif Found
Discovery/Enrichment Program
E-value
Known or Similar Motifs
MEME
8.4e-148
PAX5_HUMAN.H11MO.0.A
ZN121_HUMAN.H11MO.0.C
Reverse Complement
Show 1 More
Motif Found
Discovery/Enrichment Program
E-value
Known or Similar Motifs
MEME
1.1e-147
ZN770_HUMAN.H11MO.0.C
Reverse Complement
Show 1 More

### Supplementary Tables legends

**Supplementary Table S1. Mass spectrometry data.** (Worksheet A) Spectral count from analysis after Liquid Chromatography MS/MS of HBL-1 samples transduced with C- or N-terminally strep-tagged ETS1 using Mascot.

**Supplementary Table S2. DDX21 clinical correlation** in clinical samples**.** (Worksheet A) Genes and (Worksheet B) selected ones correlated with DDX21 in the GSE10846 dataset.

**Supplementary Table S3. Immunohistochemistry detection and analysis of DDX21 in DLBCL cases.** (Worksheet A) Intensity of staining and percentage of positive cells are reported for DDX21staining.

**Supplementary Table S4. RNASeq gene expression data after DDX21 silencing in ABC DLBCL cell lines (HBL1 and U2932).** (Worksheet A) Supervised analysis using limma of transcriptome in ABC DLBCL, (Worksheet B) U2932 and (Worksheet C) HBL1 cell lines.

**Supplementary Table S5. Small RNASeq gene expression data after DDX21 silencing in ABC DLBCL cell lines (HBL1 and U2932).** (Worksheet A) Supervised analysis using limma of miRNome, (Worksheet B) snRNome and (Worksheet C) snoRNome.

**Supplementary Table S6. DDX21 ChIP-Seq analysis in ABC DLBCL cell lines.** (Worksheet A) ChIP-Seq list of target genes. (Worksheet B) C-HiC annotation of DDX21 peaks.
